# Synaptotagmin-7 enhances phasic dopamine release

**DOI:** 10.1101/2021.10.17.464710

**Authors:** Sarah A. Kissiwaa, Joseph J. Lebowitz, Kim A. Engeln, Anna M. Bowman, John T. Williams, Skyler L. Jackman

## Abstract

Dopamine released from substantia nigra pars compacta (SNc) neurons modulates movement, motivation, and reward. In addition to their tonic firing pattern, dopamine neurons also fire high-frequency bursts that cause superlinear increases in dopamine release. To study this poorly understood form of short-term plasticity, we used the fluorescent dopamine sensor dLight1.3b to examine the role of the calcium-binding protein synaptotagmin-7 (SYT7). We report that SYT7 mediates a hidden component of facilitation, which was unmasked by lowering initial release probability, or by low-frequency stimulation of nerve terminals. In *Syt7* KO neurons, there was profound synaptic depression that significantly reduced release during stimulations that mimic *in vivo* firing patterns of SNc neurons. D2-mediated inhibitory postsynaptic currents in the SNc revealed a similar role for SYT7 in somatodendritic release. Our results indicate that SYT7 drives short-term facilitation of release from dopamine neurons, which likely underlies frequency-dependence of dopamine signaling *in vivo*.

## Introduction

Midbrain dopamine neurons modulate activity in disparate brain regions to shape affect, attention, and action (Beeler & Dreyer, 2019). Dopamine neurons fire tonically at low frequencies (∼1-8 Hz), but also fire high-frequency (>20 Hz) bursts in response to salient stimuli (Fiorillo et al., 2008; Fiorillo et al., 2003; Otomo et al., 2020). Phasic firing releases dopamine more effectively than tonic firing. For example, appetitive rewards elicit phasic firing in the midbrain (Schultz et al., 1997), and increase dopamine release in forebrain terminal regions (Day et al., 2007). Stimulating the midbrain with high-frequency bursts increases striatal dopamine concentrations more than the same number of stimuli at low frequency (Gonon, 1988; Tsai et al., 2009). These observations suggest that activity-dependent presynaptic plasticity increases dopamine release during phasic firing (Cragg, 2003). Presynaptic plasticity is generally driven by accumulation of calcium in axon terminals during high-frequency activity (Jackman & Regehr, 2017). However, the presynaptic calcium sensor that confers frequency-dependent enhancement of dopamine release has not been identified.

The vesicular Ca^2+^ sensor synaptotagmin-1 (SYT1) is required for synchronous dopamine release in the striatum (Banerjee et al., 2020). However, when *Syt1* is deleted from SNc neurons, a slow component of asynchronous striatal dopamine release remains. Such asynchronous release has been theorized to be under control of Synaptotagmin-7 (SYT7), a high-affinity Ca^2+^ sensor that mediates asynchronous release and short-term synaptic facilitation in non-dopaminergic synapses (Chen et al., 2017; Jackman et al., 2016; Luo & Südhof, 2017; Turecek & Regehr, 2018). *Syt7* RNA is expressed throughout the brain, but is enriched in SNc neurons (Saunders et al., 2018). SYT7 partially mediates release from dendrites within the SNc, and is present in axon terminals of cultured dopamine neurons (Mendez et al., 2011), but whether SYT7 regulates release from dopamine axons remains unknown. Measurement of dopamine release at sub-second intervals is required to test this hypothesis (Covey et al., 2013). Here, we used high-speed photometry of a fluorescent dopamine sensor and whole-cell electrophysiology in *Syt7* knockout mice to probe the role of SYT7 in dopamine release. Release evoked by high-frequency stimulation was reduced in *Syt7* KO animals in nerve terminals and in somatodendritic compartments. SYT7 enhanced dopamine release during transitions from tonic to phasic firing. Our data establish an important role for SYT7 in short-term plasticity and phasic dopamine release.

## Results

We immunolabeled coronal brain sections with antibodies against SYT7 and tyrosine hydroxylase (TH) (Figure 1A). In high-resolution images obtained using 3D structured illumination microscopy, SYT7-positive puncta were abundant in the striatum, both inside and outside the boundaries of TH-positive axons (Figure 1B). 3D reconstructions of SYT7 immunoreactivity within TH positive regions revealed discrete puncta throughout dopamine axons (Figure 1C-F), consistent with SYT7 localizing to the axon terminals of dopamine neurons (Mendez et al., 2011). This localization is similar to non-dopaminergic axons (Dean et al., 2012; Jackman et al., 2016; Turecek et al., 2017), suggesting a conserved function for SYT7.

**Figure 1.**
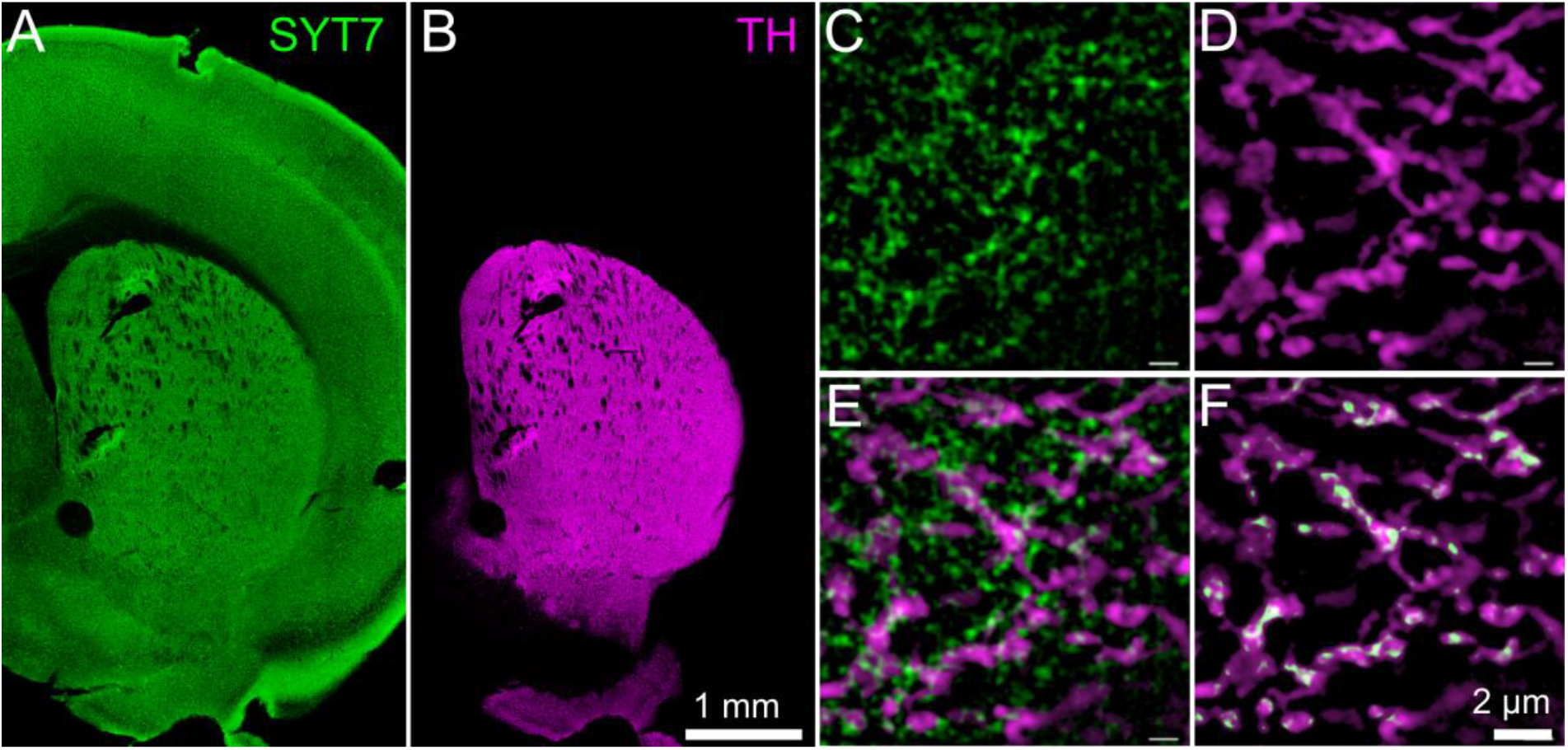
Immunohistochemistry suggests that SYT7 is present in dopaminergic axons. (**A-B**) Coronal brain sections from a WT mouse labeled with antibodies against SYT7 (A) and tyrosine hydroxylase (TH) (B). (**C-F**) 0.5 μm-thick projections of a z-stack acquired using 3D-structured illumination microscopy showing SYT7 (C) and TH labeling (D), both channels overlaid (E), and a binary mask exclusively showing SYT7-positive structures that lay within the boundaries of TH axons (F).

To probe the impact of SYT7 in striatal dopamine release, we monitored dopamine in acute striatal slices from wildtype (WT) and *Syt7*^-/-^ (*Syt7* KO) mice (Chakrabarti et al., 2003) using the genetically-encoded fluorescent dopamine sensor dLight1.3b (Patriarchi et al., 2018). Adeno-associated viral vector was injected unilaterally into the dorsal striatum to drive dLight expression in striatal neurons (Figure 2A-B). Sagittal brain slices (270 μm) were subsequently prepared for imaging and electrophysiology. dLight fluorescence was excited with dim light (mean intensity = 0.96 mW/mm^2^) and recorded at 10 kHz using a high-speed photodiode detector (Figure 2C). Dopamine axons were activated by local extracellular stimulation, in artificial cerebrospinal fluid (ACSF) containing 2 mM Ca^2+^. To isolate action potential-driven dopamine release, recordings were performed in the presence of antagonists for nicotinic, D2 and GABA receptors. Electrical stimulation elicited rapid, low noise transients (Figure 2D). The use of dim excitation light resulted in minimal photobleaching, permitting prolonged imaging over the same region of the striatum (Figure 2E). Bath application of the D1 antagonist SKF-83566 (10 μM) reduced the responses by 95 ± 3% (n = 4), confirming that fluorescence was driven by binding of dopamine to dLight (Figure 2F). Lowering external Ca^2+^ to 0.5 mM reduced dLight signals by 81 ± 3% (n = 8) consistent with the expected calcium dependence of dopamine release (Ford et al., 2010) (Figure 2G-H).

**Figure 2.**
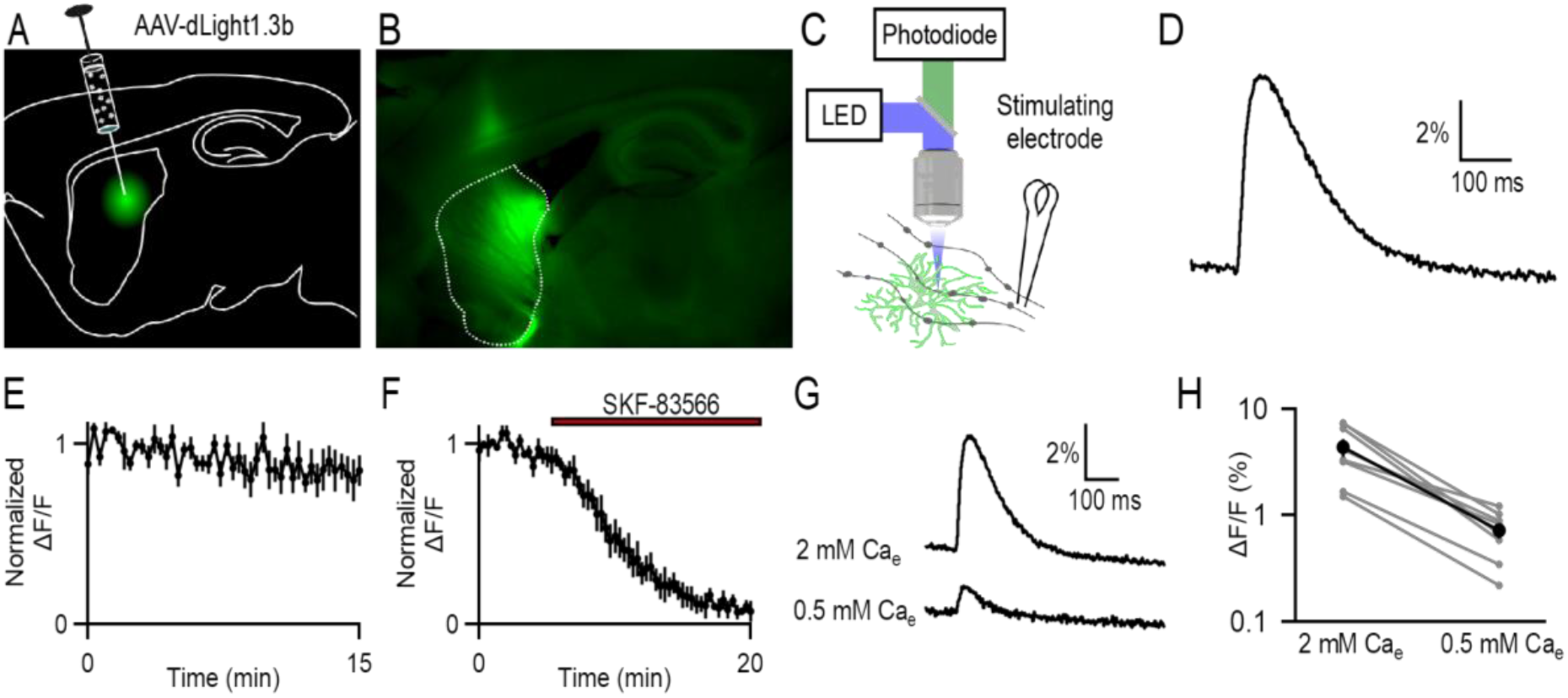
Optical monitoring of dopamine release in the striatum. (**A**) Schematic illustrating method of expressing dLight1.3b by stereotaxic injection of adeno-associated virus (AAV) into the dorsal striatum. (**B**) Representative image of a sagittal brain slice showing dLight fluorescence. (**C**) Schematic illustrating optical recording method. Fluorescence was captured using an amplified photodiode detector mounted on an upright microscope. Dopamine axons were activated by local electrical stimulation. (**D**) Representative fluorescence change evoked by electrical stimulation. Average of 10 trials. (**E**) Averaged normalized response amplitudes during repeated stimulation at 0.05 Hz (N = 7). (**F**) Average normalized responses during bath application of the selective D1 receptor antagonist SKF-83566 (N = 4). (**G**) Representative responses evoked in an acute brain slice in the presence of 2 and 0.5 mM external Ca^2+^ (Ca_e_). (**H**) ΔF/F amplitudes in high and low Ca^2+^ ACSF (N = 8). Data in all figures represents mean ± S.E.M.

dLight binds to dopamine with low affinity, and rapid on- and off-kinetics, providing an approximate measure of the time-course of extracellular dopamine. Dopamine rises and falls within ∼100 ms of a release event (Condon et al., 2021), much faster than the postsynaptic responses activated by G-protein coupled receptors. Because SYT7-mediated asynchronous release can prolong the time-course of release at other synapses (Bacaj et al., 2013; Turecek & Regehr, 2018), we evaluated whether the loss of SYT7 affected the dopamine release kinetics as measured by dLight. Neither the rise-time (20-80%), the time to the peak of the dLight transient, or the time for transients to decay to 50% of the peak were altered in the absence of SYT7 (Figure 3A-B), suggesting that SYT7 does not prolong the presence of dopamine in the extracellular space by promoting asynchronous release.

**Figure 3.**
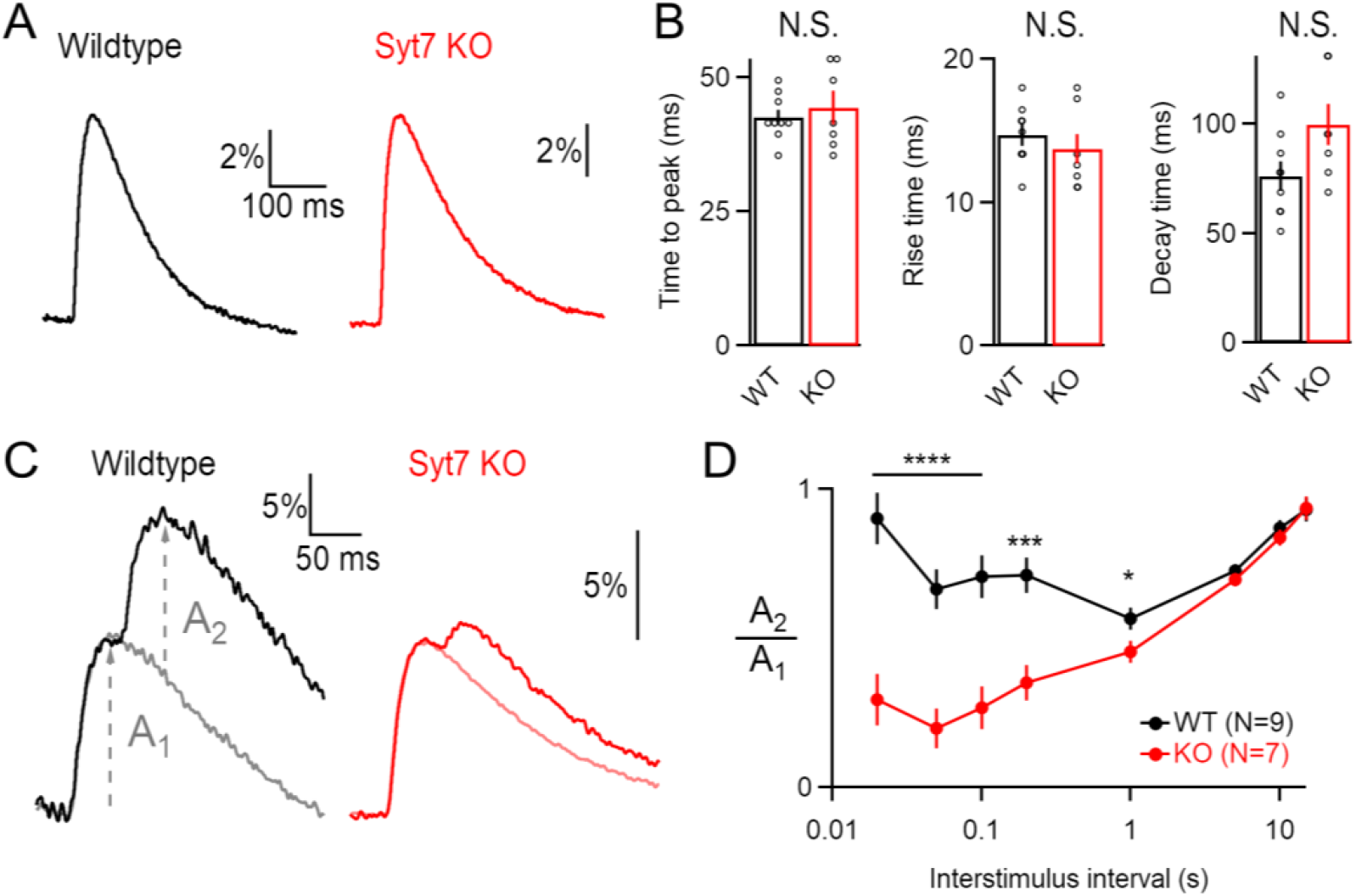
Syt7 reduces short-term depression of striatal dopamine release. (**A**) Representative dLight transients in striatal slices from WT and Syt7 KO animals. (**B**) Averaged kinetics of dLight transients. (**C**) Representative responses to single stimuli (light traces) and paired-pulse stimulation at 20 ms intervals (dark traces). (**D**) Averaged paired-pulse ratios (A_2_/A_1_) at varying interstimulus intervals. Statistical significance in all experiments was determined by unpaired Student’s t-test after normal distribution and homoscedasticity were confirmed. For comparisons of more than 2 samples, two-way ANOVA tests were performed and Šidák corrected. Significance is shown as *: p < 0.05, **: p < 0.01, ***: p < 0.001, ****: p < 0.0001.

In addition to asynchronous release, SYT7 mediates short-term synaptic facilitation. Such facilitation increases synchronous release for several hundred milliseconds after an action potential, as assessed by the paired-pulse ratio (PPR) (Jackman & Regehr, 2017). Because dLight transients lasted for more than 100 ms, for short inter-stimulus intervals (ISI) the amplitude of the second response was calculated by subtracting the response driven by a single stimulus (Figure 3C). In WT animals the PPR was ∼1 at short intervals (0.87 ± 0.07 for ISI = 20 ms), followed by marked depression, and slow recovery from depression (Figure 3D). In contrast, *Syt7* KOs displayed the hallmarks of high release probability (*p*) synapses, depressing at short intervals (PPR = 0.34 ± 0.05 for ISI = 20 ms) and recovering slowly. This suggests that SYT7 serves to offset vesicle depletion, resulting in sustained striatal dopamine release at short inter-stimulus intervals.

Synapses that show short-term depression can still possess a hidden component of facilitation that reduces depression at high stimulation frequencies (Turecek et al., 2017). This hidden facilitation can be unmasked by decreasing external Ca^2+^ (Ca_e_) to reduce *p*. However, lowering Ca_e_ to 0.5 mM decreased dLight transients to the same degree in WT and KO animals (Figure 4A-B), suggesting that the Ca^2+^-dependence of dopamine release is not affected by SYT7, as observed at other synapses (Jackman et al., 2016; Liu et al., 2014). In WT synapses, dLight responses showed a ∼2-fold facilitation that decayed within 100 ms, whereas synapses in the *Syt7* KOs exhibited mild depression (4C-D). Taken together, these experiments suggest that striatal dopamine terminals have a high initial *p* and synaptic depression, but at high firing frequencies SYT7 drives facilitation.

**Figure 4.**
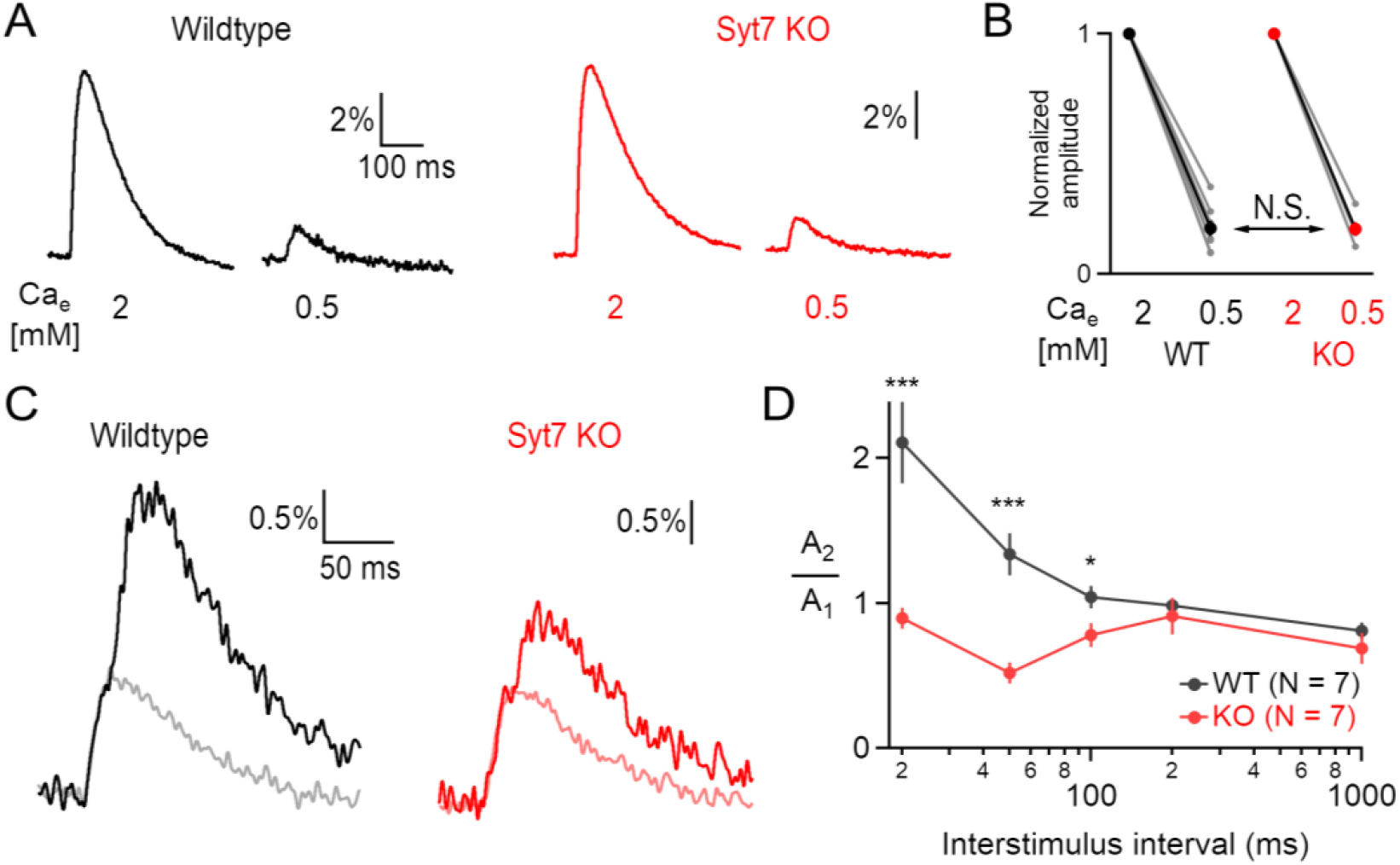
Syt7 mediates facilitation of dopamine release in low Ca_e_. (**A**) Representative dLight transients recorded in high and low Ca_e_. (**B**) Average dLight amplitudes in high and low Ca_e_, normalized to amplitudes in 2 mM Ca_e_. (**C**) Representative responses evoked by single stimuli (light) and paired-pulse stimulation at 20 ms intervals (dark) in 0.5 mM Ca_e_. (**D**) Average paired-pulse ratios in 0.5 mM Ca_e_ at varying interstimulus intervals in WT and *Syt7* KO.

Hidden facilitation can also be unmasked by depleting high-*p* vesicles with low frequency stimulation (Doussau et al., 2017; Turecek et al., 2016). This possibility is particularly relevant to dopamine release *in vivo*, as midbrain dopamine neurons switch between low-frequency tonic firing, and high-frequency phasic bursts (Schultz et al., 1997). Importantly, phasic firing is more effective at driving axonal dopamine release (Tsai et al., 2009). To explore how SYT7 affects release during physiological patterns of activity, we performed stimulations modeled after spike trains recorded *in vivo* from SNc dopamine neurons in mice (Schiemann et al., 2012). The spike pattern contained periods of tonic firing (instantaneous firing rates ranging from 1-6 Hz) as well as a tonic burst (12-33 Hz). In 2 mM Ca_e_, dopamine release was similar in WT and *Syt7* KO slices during the first 3 tonic stimuli (Figure 5A). However, at the onset of a phasic burst, dLight fluorescence increased rapidly in WT, often reaching the amplitude of the initial pre-depressed response. In contrast, in KOs release after the phasic burst was significantly decreased (Figure 5B-C). Because dopamine released during phasic bursts may accumulate and act on longer time scales, we also calculated cumulative dopamine release by integrating dLight fluorescence (Figure 5D). A significant difference in cumulative fluorescence emerged between WT and *Syt7* KO synapses after the onset of the phasic burst (Figure 5E-F). These data show that SYT7 contributes to the non-linear increase in striatal dopamine release during phasic firing of SNc neurons.

**Figure 5.**
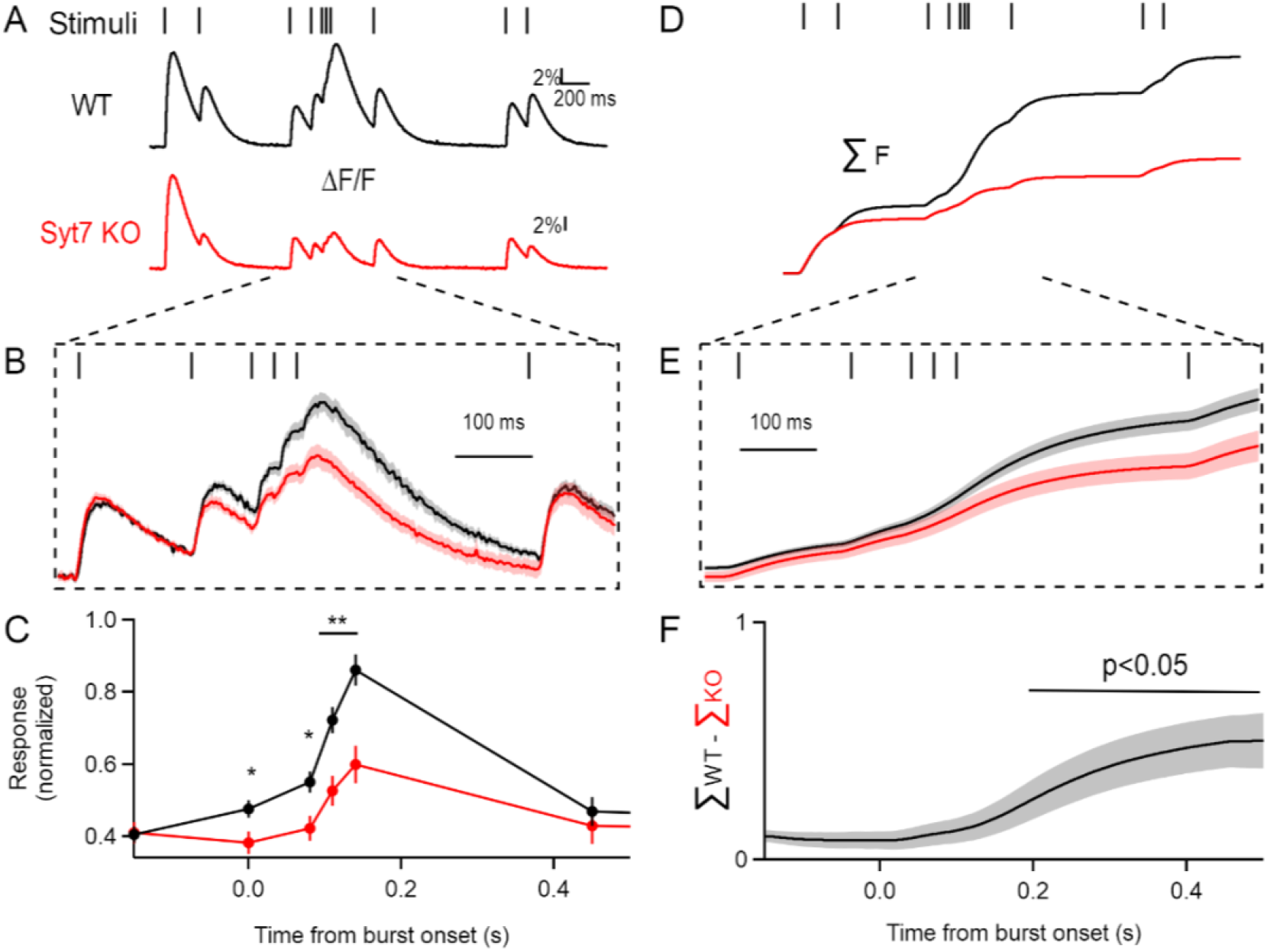
Syt7 increases dopamine release during phasic firing. (**A**) Stimulus train modeled after *in vivo* recordings from SNc dopamine neurons, and representative dLight responses from WT and *Syt7* KO animals. (**B**) Averaged responses from WT (N = 9) and KO (N = 6), normalized to the initial response amplitude. (**C**) Average normalized peak amplitudes for each stimulus, relative to the onset of the phasic burst. (**D**) Integrated dLight fluorescence (ΣF) calculated from the representative traces in (A), normalized to the end of the first response. (**E**) Averaged integrated fluorescence for WT and KO animals. (**F**) Difference in cumulative fluorescence between genotypes.

The data thus far support a role for SYT7 in driving short-term facilitation of dopamine release from striatal axon terminals during transitions from tonic to phasic firing. These transitions are in part regulated by D2 receptors expressed on dopamine cell dendrites which are activated by somatodendritic release (Lin et al., 2021). To determine the role of SYT7 in somatodendritic release, we performed whole-cell voltage-clamp recordings of D2-IPSCs. In *Syt7* KOs there was no change in the amplitudes of D2-IPSCs induced by single electrical stimuli (Figure 6A, WT: 28 pA ± 6 pA, KO: 25 ± 4 pA, p = 0.68), or a pair of stimuli at 40 Hz (Figure 6B). However, the amplitude of D2-IPSCs elicited by trains of 5 stimuli at 40Hz, normalized to the response to a single stimulus, was lower in KO animals (WT: 3.8 ± 0.4, Syt7 KO: 2.5 ± 0.2, p < 0.0001) (Figure 6C). Paired-pulse ratios for stimuli presented at 1 Hz were similar in WT and KO animals (WT: 0.65 ± 0.02, KO: 0.64 ± 0.04, p = 0.90) (Figure 6F), consistent with SYT7’s minimal contribution to release at low frequencies in other synapses (Jackman et al., 2016). To examine whether SYT7-mediated facilitation could be revealed by pre-depressing synapses with low-frequency stimulation, we stimulated the SNc with 3 pre-pulses at 1 Hz, followed by a phasic train of 5 pulses at 100 Hz. Facilitation of release was measured as the ratio of the final pre-pulse IPSC (3^rd^) and the IPSC induced by the 100 Hz phasic stimulus train (4^th^) (Beckstead et al., 2007). In WT animals, the phasic IPSC was 4.6-fold larger than the pre-depressed IPSC, a greater ratio than that seen without a series of pre-pulses. However, in *Syt7* KO slices the ratio of pre-pulse to phasic IPSC was significantly lower than in WT slices, and did not differ from that seen without pre-depression in KO slices (WT: 4.6 ± 0.2, KO: 2.6 ± 0.2, p < 0.0001) (Figure 6E). Thus, Syt7 also mediates facilitation of dopamine release at somatodendritic sites.

**Figure 6.**
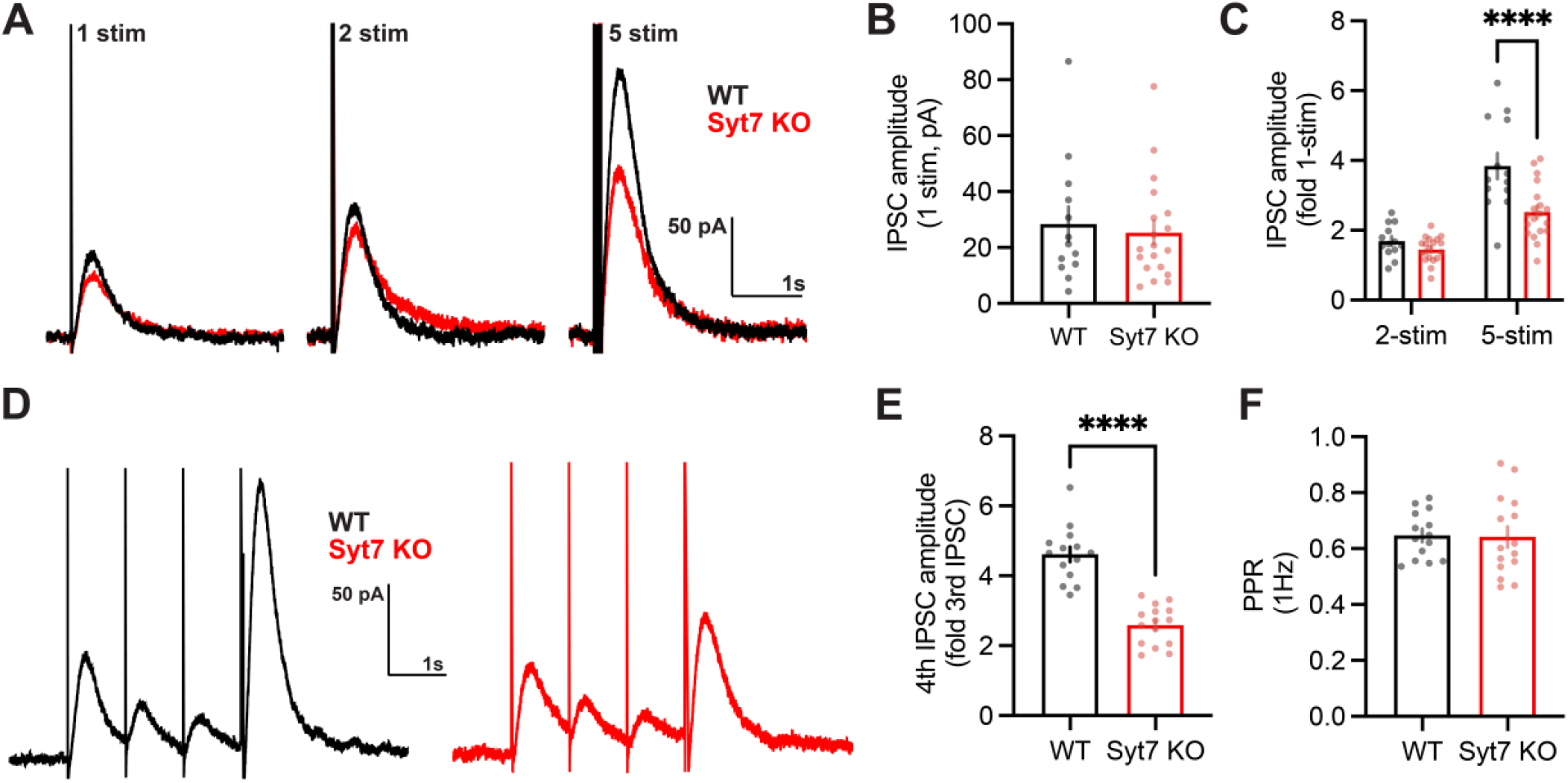
Syt7 mediates facilitation of somatodendritic dopamine release. **(A**) Representative D2-IPSCs induced by a single electrical stimulus, or by two or five stimuli at 40Hz. (**B**) Amplitude of D2-IPSCs elicited by single stimuli in WT (N = 13 cells from 4 animals) and *Syt7* KO animals (N = 19 cells from 4 animals). (**C**) Amplitude of D2-IPSC induced by two or five stimuli at 40 Hz, normalized to the response elicited with one stimulus. Same N as in (A) (**D**) Unmasking of facilitation protocol: three stimuli were presented at 1 Hz, followed by a 5-stimuli train at 100 Hz. Facilitation was quantified as the ratio of the 5-stimuli IPSC to the 3^rd^ IPSC at 1 Hz. (**E**) Quantification of facilitation in WT (N = 14 cells, 3 animals) and *Syt7* KOs (N = 15 cells, 3 animals) (**F**) PPR measured for the first two 1 Hz stimuli.

## Discussion

Here, we report that SYT7 drives short-term facilitation of dopamine release. SYT7 significantly increased release during phasic bursts, despite the fact that both axonal and dendritic synapses exhibit a high initial release probability and overt synaptic depression. A mixture of facilitation and depression was previously inferred from *in vivo* electrochemical recordings of striatal dopamine concentrations during midbrain stimulation (Montague et al., 2004). Similar “hidden” facilitation has been observed at other depressing synapses, where SYT7 maintains release during sustained firing (Chen et al., 2017; Turecek et al., 2017). Alternatively, tonic pacemaker activity *in vivo* could deplete high release probability vesicles, accentuating release during phasic bursts when SYT7 is activated by buildup of presynaptic calcium. Our experiments highlight the utility of dLight1.3b for investigating rapid dopaminergic signaling, which could be used to assess how presynaptic short-term plasticity impacts dopamine release *in vivo*.

A potential limitation of this study is the use of global *Syt7* knockout animals, which might lead to compensatory changes that confound the interpretation of our results. However, in previous studies global *Syt7* deletion abolished facilitation selectively, without changing other neuronal or synaptic properties (Bacaj et al., 2013; Jackman et al., 2016). Moreover, facilitation is restored in global knockouts in a cell-autonomous manner by presynaptic rescue of *Syt7* (Jackman et al., 2016; Turecek et al., 2017). In biochemical assays SYT7 remains bound to calcium for hundreds of milliseconds (Hui et al., 2005), similar to the time-course of facilitation we observed for dopamine release. Hence, the most parsimonious interpretation for our results is that SYT7 binds to presynaptic calcium following an action potential, and potentiates dopamine release for several hundred milliseconds.

Despite progress in understanding the molecular basis of dopamine release, exactly how neuronal activity and Ca^2+^ regulate release is unknown, and an understanding of somatodendritic release mechanisms remains particularly incomplete. Dopaminergic transmission relies on some of the same release machinery proteins that support fast, calcium-dependent neurotransmitter release from non-dopamine synapses. Dopamine release from both terminals and dendrites is dependent on RIM1, an active zone scaffolding protein involved in vesicle priming, and Ca^2+^ channel tethering (Liu et al., 2018; Robinson et al., 2019). However, dendrites and terminals may employ different Ca^2+^ sensors for release, and exhibit different Ca^2+^-dependence. The vesicular Ca^2+^ sensor SYT1 is required for release from terminals (Banerjee et al., 2020), but not for somatodendritic release (Delignat-Lavaud et al., 2021). Activity-evoked Ca^2+^ influx may not be required for release from dendrites (Chen & Rice, 2001), although this is debated (Ford et al., 2010; Yee et al., 2019). Further work is needed to clarify whether dendrites and terminals employ distinct release mechanisms. The conserved role of SYT7 in facilitating release from both compartments indicates that there are marked similarities in the molecules that underlie dendritic and terminal dopamine release.

## Methods

### Animals

All mice were handled in accordance with NIH guidelines and protocols approved by Oregon Health & Science University’s Institutional Animal Care and Use Committee. Syt7 knockout mice (Chakrabarti et al., 2003) and wild-type littermates of either sex were used. Statistical tests were not used to predetermine sample size, but sample sizes were similar to those in previous studies (Ford et al., 2010; Patriarchi et al., 2018). All electrophysiological recordings and analysis were done blind to the genotype of the animal, but for other experiments blinding and randomization were not performed.

### Immunohistochemistry

Mice were anaesthetized with ketamine/xylazine (100/10 mg/kg) and transcardially perfused with 4% paraformaldehyde (PFA) in PBS. Brains were removed and post-fixed overnight at 4 °C. 50 μm slices were prepared using a Leica VT1000S vibratome, and permeabilized for 1 h in vehicle (PBS, 10% normal goat serum, 0.3% Triton X-100), then incubated overnight at 4 °C in vehicle with primary antibodies (mouse anti-Syt7, 1:200, Neuromab N275/14; rabbit anti-TH, 1:500, Millipore AB152). After 3 washes for 30 minutes in vehicle, slices were incubated with secondary antibodies in vehicle overnight at 4 °C (Alexa flor 488 goat anti mouse, Invitrogen A11001, 1:1000; Alexa flor 546 goat anti rabbit, Invitrogen A11035, 1:500). After 3 washes for 30 minutes in PBS, slices were mounted in Prolong Gold. Images were acquired using Zeiss LSM 980 or Zeiss Elyra 7 (airyscan).

### Stereotaxic injections

Adult mice (P30-P90) were anaesthetized with ketamine/xylazine (100/10 mg/kg), placed in a stereotaxic apparatus (Kopf), and supplemented with 2% isoflurane. The head was shaved and disinfected with betadine and alcohol, an incision was made to expose the skull, and a small hole was drilled. 500 nl of AAV9.syn.dLight1.3b (4.66 10^13^ gc/ml) (Patriarchi et al., 2018) was injected unilaterally using glass pipettes (Drummond Scientific) at a rate of 100nl/minute (Nanoject 3000, Drummond). Injection coordinates from bregma were +1.4mm anterior, 1.4-1.5mm lateral and 3.5mm ventral. 10 minutes after injection the pipette was slowly retracted and scalp incision was closed with gluture. Post-injected analgesic (Carprofen, 5mg/kg) was administered subcutaneously for 48 hours.

### dLight imaging

Acute brain slices were prepared from P60-120 mice 3-6 weeks after virus injection. Animals were anesthetized with isoflurane and euthanized. Brains were quickly removed into ice-cold cutting solution containing (in mM): 125 Choline-Cl, 25 NaHCO3, 10 glucose, 2.5 KCl, 1.25 NaH_2_PO_4_, 2 Na-pyruvate, 3 (3)-myo-inositol, 4.4 ascorbic acid, 7 MgCl2, 0.5 CaCl_2_, bubbled continuously with 95% O_2_/5% CO_2_. 270 μm-thick sagittal slices were prepared using a Leica VT1200S vibratome, and transferred to a holding chamber with ACSF containing (in mM): 125 NaCl, 25 NaHCO_3_, 10 glucose, 2.5 KCl, 1.25 NaH_2_PO_4_, 2 Na-pyruvate, 3 (3)-myo-inositol, 4.4 ascorbic acid, 1 MgCl_2_, 2 CaCl_2_, bubbled continuously with 95% O_2_/5% CO_2_. Slices were stored at room temperature (24±1 °C) prior to recording.

Slices were transferred to a recording chamber and superfused with bubbled ACSF (∼2 ml/min, 34±1 °C) containing 200 μM hexamethonium, 2 μM CGP, and 0.2 μM sulpiride. A bipolar stimulating electrode was placed in the dorsal striatum, and axons were excited using a constant-voltage stimulus isolator (Digitimer DS2A). dLight fluorescence was acquired through a 60X objective on an Olympus BX51WI microscope, equipped a 488 nm LED (Thorlabs) and a FITC XF100-2 filter set (Omega). Fluorescence was captured at 10 kHz using a custom-built amplified photodiode detector. Signals were digitized with an ITC-16 (Instrutech), and acquired in Igor Pro 8 (Wavemetrics) using custom-written routines (mafPC, courtesy of Matthew Xu-Friedman).

### SNc whole-cell recordings

Acute SNc slices were prepared from P30-70 mice as previously described (Robinson et al., 2019, 2017). Animals were anesthetized with isoflurane before rapid decapitation into warmed (∼35 °C) Krebs buffer containing (in mM): 126 NaCl, 1.2 MgCl_2_, 2.4 CaCl_2_, 1.4 NaH2PO_4_, 25 NaHCO_3_, and 11 Dextrose. The brain was extracted and 222μm-thick slices were cut in the horizontal plane at room temperature with continuous bubbling of 95% O_2_/5% CO_2_. Slices were allowed to recover for 30 min at 33 °C in bubbling Krebs. Typical preparations yielded 3 slices (ventral, medial, dorsal) which were hemisected for recording experiments. Brain extraction and slicing were all performed in the presence of 10 μM MK-801 to protect from NMDA-mediated excitotoxity. Recordings were obtained from hemisected slices continuously perfused with warm (∼37 °C) Krebs containing 10 μM CNQX, 100 μM picrotoxin, and 300 nM CGP55845 to block AMPA-R, GABA-A, and GABA-B conductances respectively. Internal solution used for recording contained (in mM): 100 K-methanesulphonate, 20 NaCl, 1.5 MgCl2, 10 HEPES (K), 10 BAPTA, 2 ATP, 0.2 GTP, and 10 phosphocreatine. Giga-ohm seals were obtained using glass electrodes (1.1-1.4 MOhm resistance) and allowed to stabilize for 1-2 min before break-in. No overt difference in cell firing behavior was observed between genotypes in cell attached configuration (data not shown). Series resistance, capacitance, and membrane resistance were measured shortly after break-in using the average of 20 5 mV test pulses, and were monitored during and upon completion of all recordings. Cells were voltage-clamped at -58 mV using an AxoPatch1B amplifier and current was recorded at 1k Hz. In addition to spontaneous pacemaker activity in cell-attached configuration, DA cell identity was verified by the presence of an Ih current in response to a 50 mV hyperpolarizing step. D2-IPSCs were evoked using a platinum bipolar stimulator connected to a stimulus isolator (Warner Precision instruments) placed in the slice caudal to recorded cell bodies. All recordings are the average of 3-4 repetitions conducted once per minute that are baselined to the 500 ms immediately preceding stimulation before averaging. All recording and analysis of D2-IPSC experiments was conducted using AxoGraph software (Berkeley CA).

## Acknowledgements

We thank Stefanie Kaech Petrie, Hannah Bronstein, and the Advanced Imaging Core at the Jungers Center for microscopy assistance, and Gary Westbrook and Dennis Weingarten for helpful comments on the manuscript. This work was supported by a Whitehall Foundation Grant (SLJ), the National Institutes of Health (RO1 DA004523 to JTW), the National Institute on Drug Abuse (T32DA007262 to JJL), and the National Institute of Neurological Disorders and Stroke (P30 NS061800 to the OHSU Imaging Center).

